# The effects of gait speed on the responses to immediate and prolonged exposure to mediolateral optic flow perturbation in healthy young adults

**DOI:** 10.1101/2025.06.26.661718

**Authors:** Chunchun Wu, Tom J.W. Buurke, Rob den Otter, Claudine J.C. Lamoth, Menno P. Veldman

## Abstract

**Background:** Optic flow is vital for locomotor control and is often perturbed to study the impact of optic flow on balance control. However, it remains unclear whether gait speed influences responses to such perturbations. This study aims to examine the effects of gait speed on gait parameters following immediate and prolonged exposure to mediolateral optic flow perturbations.

**Methods:** Twenty-one young adults (23.43 ± 4.19 years) walked on an instrumented treadmill, including 3 phases: baseline (3 min), perturbation with mediolateral optic flow (8 min), and post-perturbation (3 min). Trials were conducted at 0.6, 1.2, and 1.8 m/s. Ground reaction forces and 3D motion data were collected to calculate mediolateral margin of stability (MoS), mean step length (SL), step width (SW) and their variabilities. Three repeated-measures ANOVAs (Speed by Phase) were used to compare: baseline vs. early perturbation, early vs. late perturbation, and baseline vs. post-perturbation.

**Results:** The responses to immediate and prolonged exposure to optic flow perturbation were speed dependent. Walking at slow speeds induced greater immediate responses in mediolateral gait parameters (SW and mediolateral MoS, both p < 0.001) compared to walking at faster speeds. During the perturbation phase, the adaptations were larger at faster vs. slower speeds for gait parameters in the direction of movement (SL, p = 0.007).

**Conclusion:** Immediate responses and adaptations to mediolateral optic flow perturbations are speed-dependent and larger at slower gait speeds. The responses to prolonged perturbation are interpreted as step-to-step adaptations that may inform future interventions and studies on gait speed selection.

## 1. Introduction

Visual optic flow plays a crucial role in human locomotor control. Optic flow is defined as the change in the retinal image that results from motion of the walker relative to the environment. Instantaneous visual information about the near and distant environment helps regulate locomotion at a local (step by step) and global level (direction, steering and trajectory planning (Lee et al., 1997; Patla, 1997). During everyday walking, optic flow and walking speed are coupled such that walking at, e.g., a speed of four km/hour generates an optic flow of four km/hour (Zadra & Proffitt, 2016). Gait’s dependency on optic flow information is highlighted when this information is experimentally altered as this results in immediate changes in gait parameters.

Virtual reality is commonly used to manipulate optic flow by changing the speed and/or direction of optic flow in virtual sceneries. Manipulation or removal of anterior-posterior optic flow causes young healthy individuals to adapt stride length and time to correct for any mismatch between optic flow and walking speed to satisfy the perceptual-motor coupling associated with the control of walking velocity (François et al., 2011). While changes in optic flow during forward locomotion in the mediolateral direction are less common, mediolateral balance control is critical for walking stability (Kuo & Donelan, 2010). Mediolateral stability is also essential in human bipedal gait, as walking involves a single-stance phase that requires shifts of the center of mass in the mediolateral direction. To do so, individuals adjust mediolateral step parameters such as step width (Arvin et al., 2016). Because mediolateral stability relies more on sensory information as compared to anterior-posterior balance control (O’Connor & Kuo, 2009), perturbations in the mediolateral direction are often used to study walking stability.

Manipulation of mediolateral optic flow disrupts dynamic balance during walking. Specifically, at the onset of mediolateral optic flow perturbation (comparing baseline vs. early optic flow perturbation), young adults immediately increased their step width, step length, and variability of step width (SW_variability_) and step length (SL_variability_) (Shelton et al., 2024; Thompson & Franz, 2017). The increases in step width and step length, along with their variabilities in response to the optic flow perturbation, indicate a response to prevent a decrease in gait stability. This is further evidenced by the increases in the mean mediolateral margin of stability (MoS), a measure of dynamic stability (Hof et al., 2005), and its variability (Franz et al., 2015; Thompson & Franz, 2017). When individuals are exposed to the perturbation for a longer period (>6 minutes), individuals gradually adapt their gait (comparing early vs. late optic flow perturbation); spatiotemporal gait parameters return to values observed before the introduction of the mediolateral optic flow perturbation, while the mediolateral MoS remained elevated. These ‘late’ adaptations have previously been suggested to indicate a shift from a generalized, non-task-specific, anticipatory balance control strategy to a reactive strategy using step-to-step adaptations (e.g., (Thompson & Franz, 2017)). Following the removal of the mediolateral optic flow perturbation, gait parameters return to baseline values (Richards et al., 2019; Thompson & Franz, 2017).

The impact of mediolateral optic flow perturbations has been primarily studied when participants walked on a treadmill at their preferred or a fixed speed. However, in self-paced (vs. fixed speed) experiments, individuals reduce their walking speed in response to optic flow perturbation, potentially to enhance gait stability (Richards et al., 2019; Shelton et al., 2024). Walking speed is known to influence various gait parameters in different ways (Hagoort et al., 2022; Shelton et al., 2024). Specifically, lower walking speed decreases step length and increases step width and its variability (Buurke et al., 2019; Fukuchi et al., 2019; Hagoort et al., 2022; McCrum et al., 2019). Moreover, lower walking speeds (0.5 - 1.4 m/s) were associated with reduced gait stability and poorer gait quality (complexity, intensity, smoothness) (Huijben et al., 2018). Because mediolateral optic flow is also associated with gait stability (Shelton et al., 2022), the impact of mediolateral optic flow perturbation may depend on walking speed. On the other hand, faster walking speeds are associated with higher optic flow speeds, which partially determine how individuals interpret flow information (Kountouriotis et al., 2016; Raffi et al., 2017). Therefore, gait speed likely influences an individual’s response to mediolateral optic flow perturbations.

The present study aims to establish the effects of three different gait speeds on responses to mediolateral optic flow perturbations by directly comparing gait parameters between four phases: baseline walking without perturbation, first minute of walking under optic flow perturbation (early perturbation), last minute of walking under optic flow perturbation (late perturbation) and post-perturbation. Specifically, we will:(1) Examine the effects of walking speed on the immediate responses to mediolateral optic flow perturbation on spatiotemporal gait parameters and the mediolateral MoS. We hypothesize that optic flow perturbation has the largest effect when walking at the slowest speed. (2) Determine if walking speeds influence how individuals adapt their spatiotemporal gait variables and mediolateral MoS in response to a prolonged mediolateral optic flow perturbation. We hypothesize that prolonged exposure to the mediolateral optic flow perturbation causes gait parameters to gradually return to baseline levels across all three speed conditions, reflecting a shift from a reactive to a proactive strategy. (3) Examine whether any aftereffects, i.e., the change in gait parameters following removal of optic flow perturbation values depends on walking speed. We hypothesized a return to baseline values independent of walking speed.

## 2. Methods

### 2.1. Participants

A total of 21 healthy young adults (11 females; age: 23.43 ± 4.19 years, height:177.19 ± 12.14 cm, weight: 72.14 ± 15.65 kg) participated in this study after providing written informed consent. Participants had no neurological, cognitive or orthopedic impairments that affected gait or balance and had no prior experience with optic flow perturbation experiments. The study protocol was approved by the Central Ethics Review Board of the University Medical Center Groningen (10543, January 11, 2023) and was conducted according to the Declaration of Helsinki (World Medical Association, 2013).

### 2.2. Experimental protocol

Participants walked on an instrumented treadmill (M-Gait, Motek Medical B.V., Amsterdam, The Netherlands). The continuous, 14-minute-long walking protocol consisted of three minutes of baseline walking, followed by eight minutes of perturbation, i.e., walking with an optic flow perturbation, and concluded with another three minutes of walking without perturbation (post-perturbation; Figure 1A). During the experiment, participants walked in a virtual hallway displayed on a 180° screen (Figure 1B). The hallway moved in the anterior-posterior direction at a speed that matched the walking speed. During the perturbation phase, a mediolateral optic flow perturbation was superimposed on the baseline situation using D-Flow software (Motek Medical, Amsterdam, The Netherlands). The perturbation signal *P* was based on previous studies (e.g., [8]) and consisted of a sum of three sinusoids (*φ* = 0, Equation 1, Figure 1C). Both phases were conducted at slow (0.6 m/s), intermediate (1.2 m/s), and fast (1.8 m/s) speeds. Each speed was normalized for leg length by multiplying the speed by the square root of leg length (Hof, 1996). The order of gait speeds was randomized between subjects. Participants were familiarized with each speed for one minute.

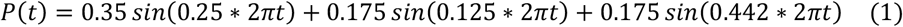

Throughout the experiment, a 10-camera motion capture system (VICON MX, Vicon Motion Systems Ltd., Oxford, UK) recorded 3D position data from markers placed on the sacrum and the heels at 100 Hz. Force plates embedded in the treadmill recorded three-dimensional ground reaction force data at 1000 Hz. Safety measures (harness and two handrails; Figure 1B), were implemented but participants were instructed not to use them.

**Figure 1.**
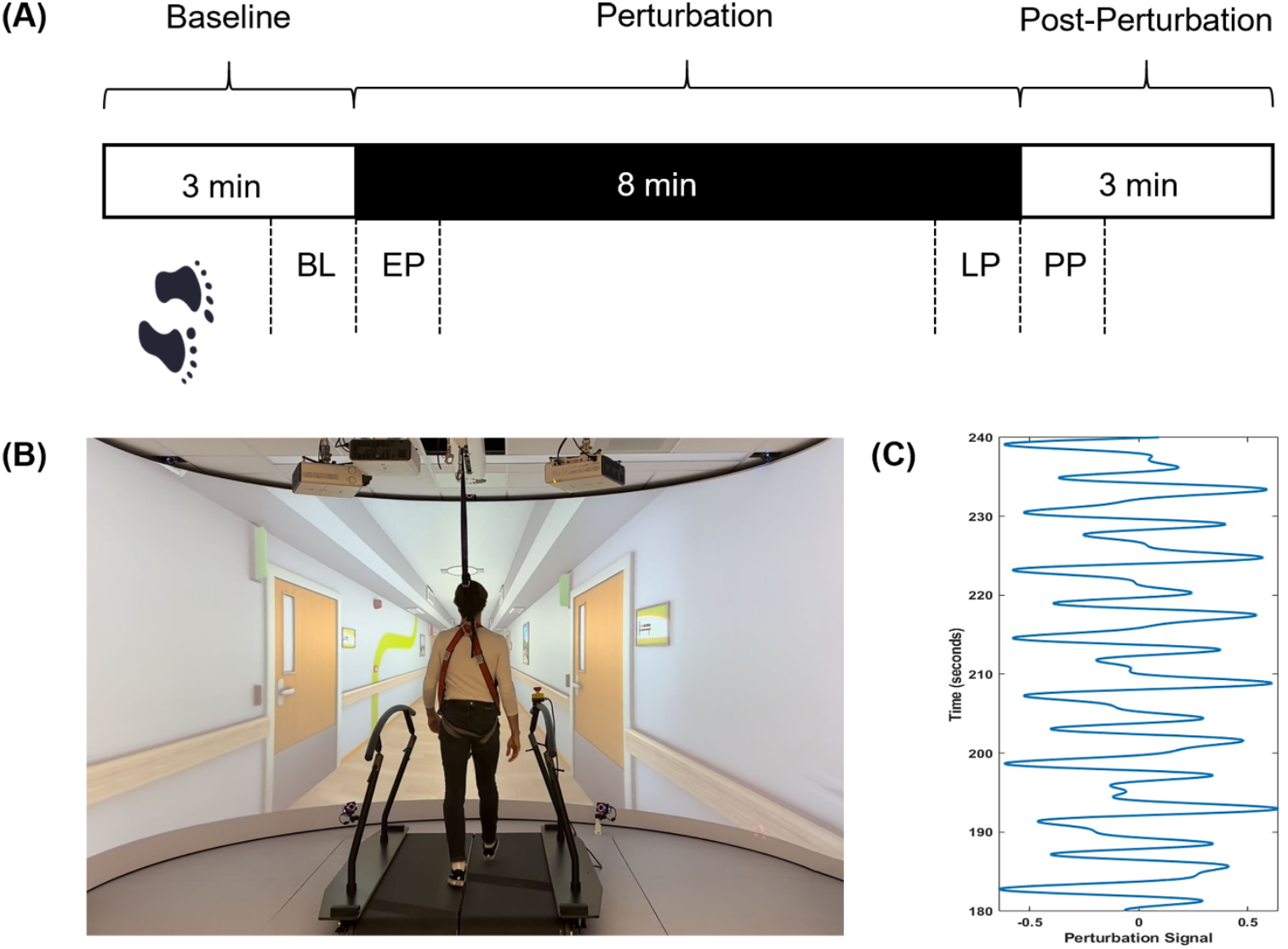
Experimental protocol. (A) Experimental protocol, with the duration of walking on the treadmill during each phase; B) Perturbation signal; (C) Participant walking on a treadmill in front of the scenery. BL: baseline; EP: early perturbation; LP: late perturbation; PP: post-perturbation

### 2.3. Data analysis

Spatiotemporal gait parameters and the mediolateral MoS were computed from the marker and force plate data, respectively, and averaged for four phases: The final 60 steps of the baseline, the first (early perturbation) and last (late perturbation) 60 steps of the perturbation phase as well as the first 60 steps of the post-perturbation phase. Marker data gaps were first fitted with a Woltring fit pipeline if gaps were shorter than 25 frames using Nexus software (Version 2.12). Then, the marker data was low pass filtered with a zero-phase Butterworth filter (2^nd^ order, 20 Hz cutoff) using custom-made software (MATLAB, vR2022B; The MathWorks Inc., Natick, MA, USA). Heel strikes and toe-offs of the right and left foot were detected by identifying the peaks and throughs, respectively, in the absolute distance between the sacrum marker and right and left heel markers (Hof, 1996). Step lengths were calculated as the anterior-posterior distance between the two heel markers at the moment of left and right heel strikes. Step widths were defined as the absolute distance in mediolateral direction between the two heel markers at the moment of two consecutive heel strikes (i.e., one left and one right heel strike). The mediolateral MoS was computed using three-dimensional ground reaction force data (Buurke et al., 2023; Curtze et al., 2024). Specifically, the extrapolated center of mass was first calculated as the vertical projection of the center of mass plus the velocity of the center of mass divided by the eigenfrequency of the effective pendulum. This eigenfrequency was calculated as the square root of the gravitational acceleration divided by leg length, quantified as the distance between the spina iliaca anterior superior to the medial malleolus of the right leg, multiplied by 1.2 (Hof et al., 2005). Given the mediolateral nature of the optic flow perturbation, we computed the mediolateral MoS in the mediolateral direction. The mediolateral MoS was defined as the minimum distance between the mediolateral Base of Support (BoS) and mediolateral extrapolated center of mass during single stance. The mediolateral BoS was defined as the mean mediolateral center of pressure position during single stance. The variability of the outcome measures, i.e., SL_variability_, SW_variability_ and mediolateral MoS_variability_, was computed as the standard deviation within each phase.

### 2.4. Statistical analysis

Statistical analyses were performed in SPSS Statistics version 28.0.1.0 (IBM). The normality of the data was tested using the Shapiro-Wilk test. To meet the assumption of normality for parametric testing, SL_variability_, SW_variability_ and mediolateral MoS_variability_ were log-transformed. To test our hypotheses, three repeated measures analyses of variance (RM ANOVA) on the mean and variability of spatiotemporal gait parameters and the mediolateral MoS were performed. First, to examine whether gait speed influenced the immediate effect of the optic flow perturbation, a three (Speed: slow, intermediate, fast) by two (Phase: baseline vs. early perturbation) RM ANOVA was performed. Second, to examine the effects of prolonged exposure to optic flow perturbation, a three (Speed) by two (Phase: early vs. late perturbation) RM ANOVA was performed. Last, to examine whether gait parameters returned to baseline levels during post-perturbation, a three (Speed) by two (Phase: Baseline vs. post-perturbation) RM ANOVA was performed. Greenhouse-Geisser correction was applied if the assumption of sphericity was violated. Partial eta-squared was used as a measure of effect size (small effect: ηp^2^ ≈ 0.01, medium effect: ηp^2^ ≈ 0.06, large effect:ηp^2^ ≈ 0.14; (Cohen, 2013)). In the case of a significant main effect of Speed or Speed by Phase interaction, paired t-tests were used to evaluate which speeds differed, using Bonferroni correction to correct for multiple comparisons. The significance threshold of all tests was set at p < 0.05.

## 3. Results

In all comparisons, step length, SW_variability_ and mediolateral MoS_variability_ were significantly increased with speed, while step width and SL_variability_ significantly decreased with faster gait speeds (Tables S1-4).

### 3.1. Immediate effects of the perturbation

The immediate effects of the optic flow perturbation (i.e., baseline vs. early perturbation) depended on gait speed. Across the three gait speed conditions, step length *(F(1,20) = 91*.*96, p < 0*.*001)* decreased and step width *(F(1,20) = 74*.*54, p < 0*.*001)*, mediolateral MoS *(F(1,20) = 58*.*18, p < 0*.*001)*, SL_variability_ *(F(1,20) = 117*.*06, p < 0*.*001)*, SW_variability_ *(F(1,20) = 156*.*48, p < 0*.*001)* and mediolateral MoS_variability_ *(F(1,20) = 139*.*41, p < 0*.*001)* increased (main effect of Phase, Table S1). Post hoc analyses following up on a significant Speed by Phase interaction (Table S1) revealed that step width and mediolateral MoS increased significantly more when walking at the slowest *(t_20_ = 3*.*46, p = 0*.*003 and t_20_ = 4*.*19, p < 0*.*001)* and intermediate gait speeds *(t_20_ = 4*.*25, p < 0*.*001 and t_20_ = 3*.*34, p = 0*.*003) speeds* compared to walking at the fast speed. Walking at an intermediate speed increased SL_variability_ significantly more than walking at a slow speed *(t_20_ = -3*.*18, p = 0*.*005; Figure 2B-D; Table S2*.*1)*.

**Figure 2.**
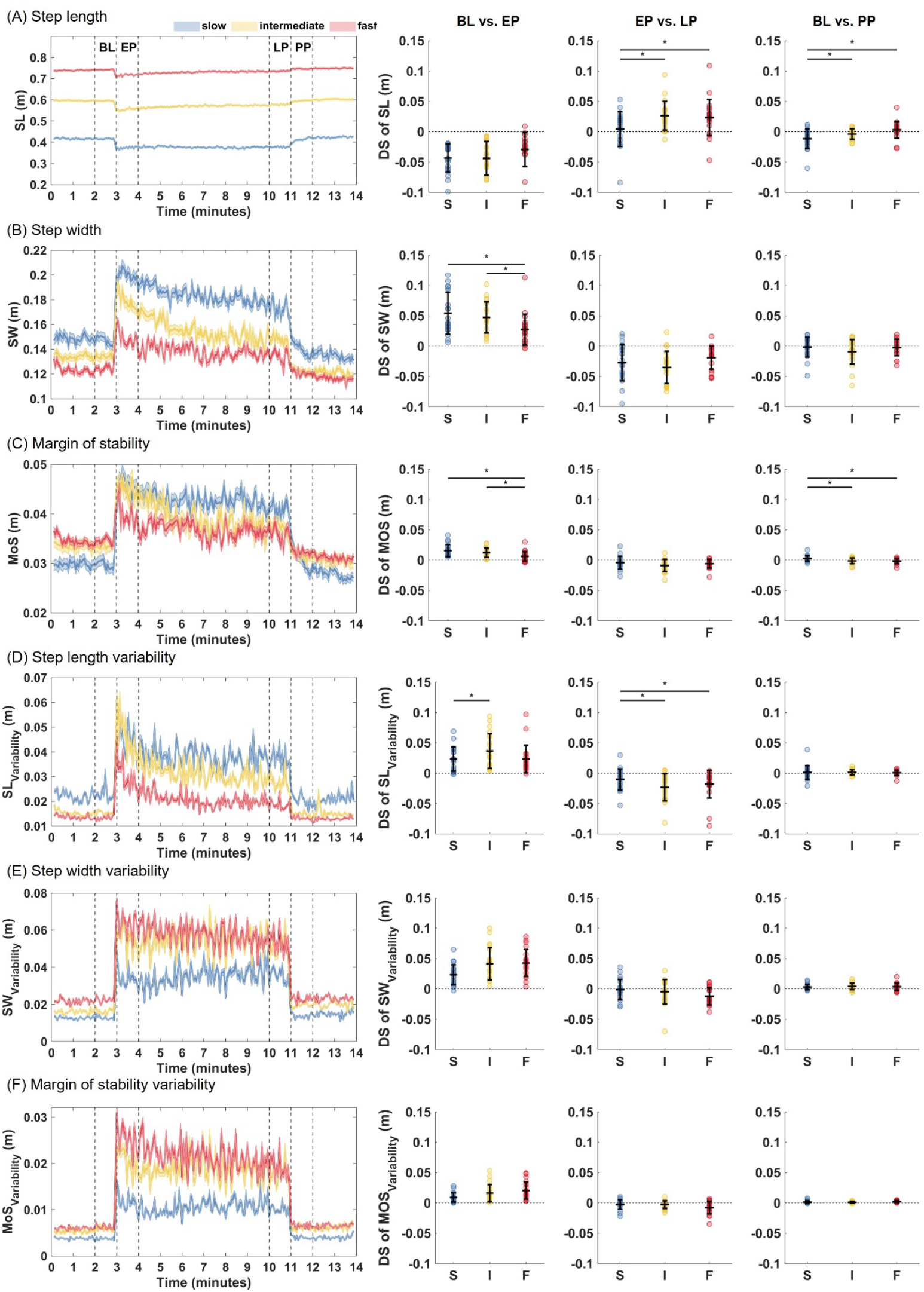
Changes in spatiotemporal gait parameters and margins of stability and their variability following immediate and prolonged optic flow perturbation. The left column of panels A-F displays the changes in gait parameters for the duration of the experimental protocol. The vertical dashed lines represent the start and end of the phases over which gait parameters are averaged for the analyses: baseline (BL), early perturbation (EP), late perturbation (LP) and post-perturbation (PP). The right three columns of panels A-F display the results of speed-by-phase interaction effects: BL vs. EP, EP vs. LP and BL vs. PP with the three different speeds (slow (S), intermediate (I) and fast(F)) on the x-axes and difference scores (DS) on the y-axes. The difference is calculated as the difference between two phases within one speed, where the dashed horizontal line indicates no change. SL: step length; SW: step width; mediolateral MoS: Margin of Stability; *: p-value < 0.05

### 3.2. Effects of prolonged exposure to the perturbation

Responses to prolonged exposure to the optic flow perturbation (early vs. late perturbation) showed a significant main effect of Phase and a Speed by Phase interaction. Across all speeds, step length *(F(1,20) = 16*.*97, p < 0*.*001)*, step width *(F(1,20) = 42*.*70, p < 0*.*001)*, mediolateral MoS *(F(1,20) = 15*.*06, p = 0*.*001)*, SL_variability_ *(F(1,20) = 42*.*26, p < 0*.*001)* and mediolateral MoS_variability_ *(F(1,20) = 7*.*59, p = 0*.*013)* returned in the direction of baseline levels (Table S3). Post hoc analyses (Table S3) showed that the changes from early to late perturbation in step length and SL_variability_ differed between gait speed conditions. Specifically, the increases and decreases in step length and SL_variability_, respectively, were lower when walking at slow speed compared to walking at intermediate *(t_20_ = -3*.*66, p = 0*.*002 and t_20_ = 2*.*63, p = 0*.*016)* and fast speeds *(t_20_ = -2*.*32, p = 0*.*031 and t_20_ = 3*.*18, p = 0*.*005; Figure 2A and 2D; Table S2*.*2)*.

### 3.3. Effects after removal of the perturbation

The extent to which gait parameters returned to baseline levels when the optic flow perturbation was removed, differed between gait speed conditions. The main effect of Phase showed that step length *(F(1,20) = 4*.*74, p = 0*.*042)* was decreased and SW_variability_ *(F(1,20) = 25*.*81, p < 0*.*001)* and mediolateral MoS_variability_ *(F(1,20) = 39*.*30, p < 0*.*001)* were increased at post perturbation as compared to baseline (Table S4). Moreover, the results showed a significant Speed by Phase interaction (Table S4). Post hoc analyses revealed that the decrease and increase in step length and mediolateral MoS, respectively, was greater when walking at the slow speed as compared to intermediate *(t_20_ = -2*.*39, p = 0*.*027 and t_20_ = 3*.*2, p = 0*.*005)* and fast speeds *(t_20_ = - 3*.*34, p = 0*.*003 and t_20_ = 3*.*28, p = 0*.*004; Figures 2A and 2C; Table S2*.*3)*.

## 4. Discussion

This study examined the effect of gait speed on the immediate and prolonged responses to mediolateral optic flow perturbation as well as the effects during post-perturbation on gait parameters. The results revealed speed dependent immediate effects of the mediolateral optic flow perturbation on gait parameters in the frontal plane, indicated by higher values for step width and mediolateral MoS when walking at slow vs. faster speeds. Following prolonged exposure to the optic flow perturbation, these parameters gradually returned in the direction of baseline levels, but their variability did not. In addition, changes in step length and SL_variability_ were less pronounced at slow vs. faster speeds following prolonged exposure to optic flow perturbation. Following the removal of the perturbation, step width and mediolateral MoS returned to baseline levels, but their variability remained high. Altogether, the data indicate that step-to-step adaptations in response to mediolateral optic flow perturbation depend on gait speed.

Optic flow perturbation caused speed-dependent changes in gait parameters immediately following the onset of the exposure. In agreement with previous studies, challenging walking balance with mediolateral optic flow perturbation caused individuals to take shorter and wider steps (Richards et al., 2019; Salinas et al., 2017; Shelton et al., 2024; Thompson & Franz, 2017), and increase the variability of gait parameters (Thompson & Franz, 2017). Our data further confirm previously observed trends for increases in mediolateral MoS (Richards et al., 2019). Interestingly, the current data revealed, for the first time in the context of optic flow perturbation, that the changes in frontal plane gait parameters (step width and mediolateral MoS), were more pronounced at slower gait speeds while their variability, indicative of reactive balance control (O’Connor et al., 2012; O’Connor & Kuo, 2009), remained low. Walking at slow speed is inherently more unstable than walking at faster speeds (Buurke et al., 2019), is associated with a lower step frequency (Ziegler et al., 2024) and the higher single support time present at slower walking speeds leads to a smaller mediolateral MoS (Buurke et al., 2019). Provided that lateral foot placement relies more on the integration of information from various sensory sources than foot placement in the direction of movement (Bauby & Kuo, 2000; Collins & Kuo, 2013; Donelan et al., 2004; O’Connor et al., 2012; O’Connor & Kuo, 2009), the speed-dependent effects on step width and mediolateral MoS might imply that the disintegration of sensory information caused by mediolateral optic flow perturbation is larger at slower gait speeds. This is supported by findings that the Central nervous system relies more on vestibular input at slower speeds, making sensory perturbations more disruptive during slow walking (Brandt et al., 1999). The lower variability in step width and mediolateral MoS at slow vs. faster walking speeds may even suggest that optic flow perturbation affects anticipatory balance control. This suggestion is in line with findings that variability in gait parameters only increased once gait parameters returned to baseline values following prolonged exposure to mediolateral optic flow perturbation (Thompson & Franz, 2017).

The responses to prolonged exposure to optic flow perturbation were also speed-dependent. Consistent with earlier results (Thompson & Franz, 2017), all gait parameters except for the variability in step width returned in the direction of baseline levels. In contrast with the immediate effects, the time-dependent adaptations differed between speeds for step length and its variability and not for mediolateral parameters. Since step length relies less on the integration of various sensory sources (Bauby & Kuo, 2000; Collins & Kuo, 2013; Donelan et al., 2004; O’Connor et al., 2012; O’Connor & Kuo, 2009), step-to-step adaptations in response to optic flow perturbation are possibly easier to perform for step length than for mediolateral parameters. If true, the fewer opportunities to perform step-to-step adaptations during slow vs. faster speeds, caused by reduced step frequencies (Huijben et al., 2018), may explain the speed-dependent effects for step length but not step width and mediolateral MoS.

The observation across studies that participants tend to adjust their walking pattern towards their baseline walking is consistent with the concept of sensory reweighting and might suggest that individuals prioritize maintaining balance but return to an energetically efficient walking pattern as soon as possible. Specifically, perturbation of optic flow may have triggered a shift from using unreliable visual information to reliable somatosensory information (Mahboobin et al., 2005). This shift may have supported the gradual return of step width, mediolateral MoS and mediolateral MoS_variability_ to baseline levels, leading to increasingly energetically efficient walking over the course of the perturbation phase (Kuo et al., 2005; O’Connor et al., 2012).

Once the optic flow perturbation was removed, step width and mediolateral MoS_variability_ returned to baseline levels in line with prior studies (Richards et al., 2019; Thompson & Franz, 2017). Inconsistent with the literature, this was not the case for step length and mediolateral MoS (Richards et al., 2019; Thompson & Franz, 2017). This deviation may be explained by the greater instability (Beauchet et al., 2009) and attentional demands associated with slow walking (Lajoie et al., 2016), which can delay motor responses and hinder the recovery of step length and mediolateral MoS following the removal of optic flow. Moreover, the present data did not show the effects typically observed after removal of unilateral gait perturbations (e.g., split-belt walking (Buurke et al., 2018), where responses often reverse direction as a result of motor recalibration (Mahboobin et al., 2005). The absence of these effects in response to optic flow perturbation indicates that optic flow may not cause recalibration of motor control but rather triggers step-to-step adaptations. This discrepancy may be due to the unilateral nature of split-belt perturbations, which primarily affect symmetry-based measures and elicit after-effects (Buurke et al., 2018), whereas the bilateral nature of optic flow perturbations does not disrupt symmetry and may therefore prevent recalibration. Follow-up studies adopting unilateral optic flow perturbations are needed to confirm this hypothesis. Alternatively, it is possible that optic flow perturbation does not cause a discrepancy between control signals and predicted sensory consequences but rather causes conflicting incoming sensory information that is (partly) resolved by reweighting the sensory information triggering the step-by-step adaptations.

There are several limitations of this study. The continuous perturbation signal used in this study consisted of three sinusoids of different frequencies (Thompson & Franz, 2017). Although challenging, it is possible that participants made predictions about the perturbation that obscured some effects of optic flow perturbation. Moreover, we did not control for strategies to cope with the perturbation and whether the participants kept their eyes open, although we instructed the participants to look straight to the virtual hallway. Lastly, because only healthy young individuals were included in the present study, the current results cannot be generalized to older or clinical populations. As suggested in a previous paper (Qiao et al., 2019), future studies are needed to explore the potential application of optic flow perturbation in a rehabilitation context. In line with this, walking on a treadmill access to the peripheral field of view is limited as compared to walking in physical space. To overcome this limitation, future studies could consider including eye-tracking to control for any fixations or saccades during optic flow perturbation experiments.

## 5. Conclusion

The present study showed that the immediate responses to mediolateral optic flow perturbation and the responses to prolonged exposure are amplified when walking at slow speed compared to walking at faster speeds. The changes in mediolateral and anterior-posterior gait parameters in response to optic flow perturbation are suggested to be mediated by the importance and availability of sensory information for the mediolateral and anterior-posterior gait parameters. Altogether, the data indicate that the step-to-step adaptations in response to mediolateral optic flow perturbation depend on gait speed.

## CRediT authorship contribution statement

**Chunchun Wu:** Writing – review & editing, Writing – original draft, Investigation, Visualization, Methodology, Conceptualization, Data analysis. **Tom J.W. Buurke:** Writing – review & editing, Visualization, Methodology, Conceptualization, Data analysis. **Rob den Otter:** Writing – review & editing, Methodology, Conceptualization. **Claudine J.C. Lamoth:** Writing – review & editing, Writing – original draft, Methodology, Conceptualization. **Menno P. Veldman:** Writing – review & editing, Writing – original draft, Visualization, Methodology, Conceptualization, Data analysis.

## Funding and acknowledgement

This work was supported by China Scholarship Council – University of Groningen Scholarship (C. Wu: 202206320033) and Research Foundation - Flanders (FWO; T.J.W. Buurke: 12ZJ922N). The authors declare no conflicts of interest.

## Data availability

Due to data protection, the datasets collected and analyzed during this study are not openly available. However, reasonable requests for data supporting the findings of this study are available from the corresponding author on reasonable request.

## Notes

### Competing Interest Statement

The authors have declared no competing interest.

