## Supplementary tables for "The effects of gait speed on the responses to immediate and prolonged exposure to mediolateral optic flow perturbation in healthy young adults"

**Supplementary materials**

| Table S1: Effects of optic flow perturbation and speed on gait parameters compared to BL and EP | | | | | | | | Statistics | | | |
| --- | --- | --- | --- | --- | --- | --- | --- | --- | --- | --- | --- |
|  | Slow speed | | Intermediate speed | | Fast speed | | Effect | | F_(df)_ | *p* | η_p_^2^ |
|  | BL | EP | BL | EP | BL | EP |  |  |  |  |  |
| SL | 41.64±4.70 | 37.30±4.75 | 59.59±4.73 | 55.20±4.90 | 74.28±4.49 | 71.35±4.16 | Speed | | F_(2,40)_=788.71 | <0.001 | 0.975 |
|  |  |  |  |  |  |  | Phase | | F_(1,20)_=91.96 | <0.001 | 0.821 |
|  |  |  |  |  |  |  | Speed*Phase | | F_(2,40)_=2.74 | 0.077 | 0.120 |
| SW | 14.65±4.85 | 20.07±6.07 | 13.46±4.42 | 18.18±5.71 | 12.54±3.02 | 15.25±3.93 | Speed | | F_(2,40)_=12.44 | <0.001 | 0.383 |
|  |  |  |  |  |  |  | Phase | | F_(1,20)_=74.54 | <0.001 | 0.788 |
|  |  |  |  |  |  |  | Speed*Phase | | F_(2,40)_=8.82 | <0.001 | 0.306 |
| MoS | 2.96±1.40 | 4.52±2.10 | 3.60±1.19 | 4.85±1.75 | 3.48±0.94 | 4.10±1.25 | Speed | | F_(2,38)_=2.12 | 0.134 | 0.100 |
|  |  |  |  |  |  |  | Phase | | F_(1,19)_=58.18 | <0.001 | 0.754 |
|  |  |  |  |  |  |  | Speed*Phase | | F_(2,38)_=11.66 | <0.001 | 0.380 |
| SL_var_ | 2.57±0.99 | 4.89±1.98 | 1.60±0.28 | 5.28±2.93 | 1.45±0.42 | 3.78±2.45 | Speed | | F_(2,40)_=16.64 | <0.001 | 0.454 |
|  |  |  |  |  |  |  | Phase | | F_(1,20)_=117.06 | <0.001 | 0.854 |
|  |  |  |  |  |  |  | Speed*Phase | | F_(2,40)_=5.02 | 0.011 | 0.201 |
| SW_var_ | 1.58±0.25 | 3.91±1.68 | 1.87±0.41 | 6.01±2.63 | 2.34±0.43 | 6.61±2.10 | Speed | | F_(2,40)_=48.49 | <0.001 | 0.708 |
|  |  |  |  |  |  |  | Phase | | F_(1,20)_=156.48 | <0.001 | 0.887 |
|  |  |  |  |  |  |  | Speed*Phase | | F_(2,40)_=3.17 | 0.053 | 0.137 |
| MoS_Var_ | 0.42±0.09 | 1.33±0.78 | 0.49±0.09 | 2.10±1.40 | 0.63±0.13 | 2.66±1.35 | Speed | | F_(2,38)_=34.45 | <0.001 | 0.645 |
|  |  |  |  |  |  |  | Phase | | F_(1,19)_=139.41 | <0.001 | 0.880 |
|  |  |  |  |  |  |  | Speed*Phase | | F_(2,38)_=2.85 | 0.070 | 0.131 |

All data are presented as mean ± SD in centimeters. BL: baseline, EP: early perturbation, SL: step length, SW: step width, MoS: mediolateral margin of stability, SL_var_: step length variability, SW_var_: step width variability, MoS_var_: mediolateral margin of stability variability.

| Table S2: Post hoc test for interaction effect and main effect of speed found in the RM ANOVA | | | | | | | | | | | | | | | |
| --- | --- | --- | --- | --- | --- | --- | --- | --- | --- | --- | --- | --- | --- | --- | --- |
| S2.1 Effects of optic flow perturbation and speed on gait parameters from BL to EP | | | | | | | | statistics | | | | | | | |
|  | |  | Slow | Intermediate | Fast | Pair 1  (Slow vs. Intermediate) | | | | Pair 2  (Slow vs. Fast) | | | Pair 3  (Intermediate vs. Fast) | | |
|  |  |  |  |  |  | t_df_ | *p* | | change (%) | t_df_ | *p* | change (%) | t_df_ | *p* | change (%) |
| BL  vs. EP | Main effect of speed | SL | 39.47±5.16 | 57.40±5.25 | 72.81±4.52 | t_20_=-22.61 | <0.001 | | 45.4 | t_20_=-33.20 | <0.001 | 84.5 | t_20_=-22.23 | <0.001 | 26.9 |
|  |  | SW | 17.36±6.08 | 15.82±5.58 | 13.89±3.72 | t_20_=2.57 | 0.018 | | -8.9 | t_20_=4.43 | <0.001 | -20.0 | t_20_=2.77 | 0.012 | -12.2 |
|  |  | SL_var_ | 3.73±1.94 | 3.44±2.77 | 2.61±2.10 | t_20_=3.41 | 0.003 | | -7.8 | t_20_=6.02 | <0.001 | -29.9 | t_20_=2.48 | 0.022 | -24.0 |
|  |  | SW_var_ | 2.74±1.67 | 3.94±2.80 | 4.48±2.63 | t_20_=-6.07 | <0.001 | | 43.7 | t_20_=-11.34 | <0.001 | 63.3 | t_20_=-3.33 | 0.003 | 13.6 |
|  |  | MoS_var_ | 0.88±0.72 | 1.29±1.27 | 1.65±1.40 | t_19_=-3.78 | 0.001 | | 47.3 | t_20_=-8.87 | <0.001 | 87.7 | t_19_=-4.55 | <0.001 | 27.4 |
|  | Interaction effect | SW | 5.42±3.46 | 4.73±2.55 | 2.71±2.55 | t_20_=0.98 | 0.341 | | -13.0 | t_20_=3.46 | 0.003 | -50.0 | t_20_=4.25 | <0.001 | -42.6 |
|  |  | MoS | 1.56±0.99 | 1.18±0.79 | 0.63±0.77 | t_19_=1.91 | 0.072 | | -25.0 | t_20_=4.19 | <0.001 | -62.5 | t_19_=3.34 | 0.003 | -50.0 |
|  |  | SL_var_ | 2.32±2.01 | 3.68±2.86 | 2.33±2.30 | t_20_=-3.18 | 0.005 | | 60.9 | t_20_=-0.80 | 0.087 | 0.0 | t_20_=1.41 | 0.175 | -37.8 |
| S2. 2: Effects of optic flow perturbation and speed on gait parameters from EP to LP | | | | | | | |  | | | | | | | |
| EP  vs. LP | Main effect of speed | SL | 37.53±4.62 | 56.52±4.89 | 72.52±4.76 | t_20_=-22.90 | <0.001 | | 50.6 | t_20_=-34.86 | <0.001 | 93.3 | t_20_=-20.42 | <0.001 | 28.3 |
|  |  | SW | 18.70±5.58 | 16.43±5.29 | 14.29±3.73 | t_20_=3.15 | 0.005 | | -12.1 | t_20_=4.91 | <0.001 | -23.6 | t_20_=3.23 | 0.004 | -13.0 |
|  |  | SL_var_ | 4.38±1.81 | 4.11±2.49 | 2.88±1.98 | t_20_=1.53 | 0.141 | | -6.1 | t_20_=5.91 | <0.001 | -34.3 | t_20_=3.24 | 0.004 | -30.0 |
|  |  | SW_var_ | 3.85±1.75 | 5.78±2.56 | 6.00±1.90 | t_20_=-7.38 | <0.001 | | 50.2 | t_20_=-9.00 | <0.001 | 56.0 | t_20_=-1.22 | 0.236 | 3.8 |
|  |  | MoS_var_ | 1.20±0.68 | 1.97±1.35 | 2.29±1.22 | t_19_=-4.09 | <0.001 | | 63.9 | t_20_=-8.07 | <0.001 | 90.8 | t_19_=-2.14 | 0.045 | 16.4 |
|  | Interaction effect | SL | 0.44±2.88 | 2.64±2.38 | 2.34±2.98 | t_20_=-3.66 | 0.002 | | 550.0 | t_20_=-2.32 | 0.031 | 475.0 | t_20_=0.42 | 0.680 | -11.5 |
|  |  | SL_var_ | -1.03±1.75 | -2.33±2.23 | -1.81±2.28 | t_20_=2.63 | 0.016 | | 130.0 | t_20_=3.18 | 0.005 | 80.0 | t_20_=0.47 | 0.643 | -21.7 |
| S2. 3: Effects of optic flow perturbation and speed on gait parameters from BL to PP | | | | | | | |  | | | | | | | |
| BL  vs. PP | Main effect of speed | SL | 41.06±4.89 | 59.40±4.66 | 74.43±4.64 | t_20_=-23.24 | <0.001 | | 44.7 | t_20_=-35.57 | <0.001 | 81.3 | t_20_=-30.92 | <0.001 | 25.3 |
|  |  | SW | 14.56±4.65 | 12.98±4.04 | 12.42±2.75 | t_20_=3.28 | 0.004 | | -10.9 | t_20_=3.13 | 0.005 | -14.8 | t_20_=1.10 | 0.287 | -4.3 |
|  |  | SL_var_ | 2.62±0.91 | 1.68±0.38 | 1.48±0.38 | t_20_=8.37 | <0.001 | | -35.8 | t_20_=8.42 | <0.001 | -43.4 | t_20_=2.97 | 0.008 | -11.8 |
|  |  | SW_var_ | 1.73±0.46 | 2.08±0.50 | 2.52±0.55 | t_20_=-3.79 | 0.001 | | 20.2 | t_20_=-7.65 | <0.001 | 45.6 | t_20_=-5.02 | <0.001 | 21.2 |
|  |  | MoS_var_ | 0.50±0.18 | 0.54±0.14 | 0.74±0.19 | t_19_=-2.91 | 0.009 | | 8.9 | t_20_=-10.63 | <0.001 | 48.1 | t_19_=-9.78 | <0.001 | 36.1 |
|  | Interaction effect | SL | -1.15±1.61 | -0.38±0.86 | 0.31±1.38 | t_20_=-2.39 | 0.027 | | -66.7 | t_20_=-3.34 | 0.003 | -125.0 | t_20_=-1.89 | 0.073 | -175.0 |
|  |  | MoS | 0.31±0.44 | -0.10±0.46 | -0.16±0.39 | t_19_=3.20 | 0.005 | | -133.3 | t_20_=3.28 | 0.004 | -166.7 | t_19_=0.58 | 0.569 | 100.0 |

All data are presented as mean ± SD in centimeters. BL: baseline, EP: early perturbation, LP: late perturbation PP: post perturbation, SL: step length, SW: step width, MoS: mediolateral margin of stability, SL_var_: step length variability, SW_var_: step width variability, MoS_var_: mediolateral margin of stability variability.

| Table S3: Effects of optic flow perturbation and speed on gait parameters compared to EP and LP | | | | | | | | | | | | | | Statistics | | | | | | |
| --- | --- | --- | --- | --- | --- | --- | --- | --- | --- | --- | --- | --- | --- | --- | --- | --- | --- | --- | --- | --- |
|  | Slow speed | | | Intermediate speed | | | | Fast speed | | | | Effect | | | F (df) | | *p* | | η_p_^2^ | |
|  | EP | LP | | EP | | LP | | EP | | LP | |  | | |  | |  | |  | |
| SL | 37.30±4.75 | | 37.75±4.58 | | 55.20±4.90 | | 57.84±4.61 | | 71.35±4.16 | | 73.69±5.13 | | Speed | | | F_(2,40)_=797.39 | | <0.001 | | 0.976 |
|  |  |  |  |  |  |  |  |  |  |  |  |  | Phase | | | F_(1,20)_=16.97 | | <0.001 | | 0.459 |
|  |  |  |  |  |  |  |  |  |  |  |  |  | Speed*Phase | | | F_(2,40)_=5.60 | | 0.007 | | 0.219 |
| SW | 20.07±6.07 | | 17.33±4.80 | | 18.18±5.71 | | 14.68±4.27 | | 15.25±3.93 | | 13.34±3.34 | | Speed | | | F_(2,40)_=16.53 | | <0.001 | | 0.452 |
|  |  |  |  |  |  |  |  |  |  |  |  |  | Phase | | | F_(1,20)_=42.70 | | <0.001 | | 0.681 |
|  |  |  |  |  |  |  |  |  |  |  |  |  | Speed*Phase | | | F_(2,40)_=3.03 | | 0.059 | | 0.132 |
| MoS | 4.52±2.10 | | 4.12±1.58 | | 4.85±1.75 | | 3.96±1.39 | | 4.10±1.25 | | 3.53±1.32 | | Speed | | | F_(2,38)_=3.23 | | 0.051 | | 0.145 |
|  |  |  |  |  |  |  |  |  |  |  |  |  | Phase | | | F_(1,19)_=15.06 | | 0.001 | | 0.442 |
|  |  |  |  |  |  |  |  |  |  |  |  |  | Speed*Phase | | | F_(2,38)_=2.11 | | 0.135 | | 0.100 |
| SL_var_ | 4.89±1.98 | | 3.86±1.50 | | 5.28±2.93 | | 2.95±1.14 | | 3.78±2.45 | | 1.97±0.53 | | Speed | | | F_(2,40)_=14.36 | | <0.001 | | 0.418 |
|  |  |  |  |  |  |  |  |  |  |  |  |  | Phase | | | F_(1,20)_=42.26 | | <0.001 | | 0.679 |
|  |  |  |  |  |  |  |  |  |  |  |  |  | Speed*Phase | | | F_(2,40)_=5.15 | | 0.010 | | 0.205 |
| SW_var_ | 3.91±1.68 | | 3.78±1.87 | | 6.01±2.63 | | 5.54±2.53 | | 6.61±2.10 | | 5.39±1.50 | | Speed | | | F_(2,40)_=39.77 | | <0.001 | | 0.665 |
|  |  |  |  |  |  |  |  |  |  |  |  |  | Phase | | | F_(1,20)_=3.61 | | 0.072 | | 0.153 |
|  |  |  |  |  |  |  |  |  |  |  |  |  | Speed*Phase | | | F_(2,40)_=1.48 | | 0.241 | | 0.069 |
| MoS_var_ | 1.33±0.78 | | 1.07±0.55 | | 2.10±1.40 | | 1.84±1.33 | | 2.66±1.35 | | 1.92±0.95 | | Speed | | | F_(1.88,35.71)_=22.63 | | <0.001 | | 0.544 |
|  |  |  | |  | |  | |  | |  | | Phase | | | F_(1,19)_=7.59 | | 0.013 | | 0.286 | |
|  |  |  |  |  |  |  |  |  |  |  |  | Speed*Phase | | | F_(1.55,29.48)_=1.48 | | 0.242 | | 0.072 | |

All data are presented as mean ± SD in centimeters. EP: early perturbation, LP: late perturbation, SL: step length, SW: step width, MoS: mediolateral margin of stability, SL_var_: step length variability, SW_var_: step width variability, MoS_var_: mediolateral margin of stability variability.

| Table S4: Effects of optic flow perturbation and speed on gait parameters compared to BL and PP | | | | | | | | Statistics | | | |
| --- | --- | --- | --- | --- | --- | --- | --- | --- | --- | --- | --- |
|  | Slow speed | | Intermediate speed | | Fast speed | | Effect | | F | *p* | η_p_^2^ |
|  | BL | PP | BL | PP | BL | PP |  |  |  |  |  |
| SL | 41.64±4.70 | 40.48±5.12 | 59.59±4.73 | 59.21±4.71 | 74.28±4.49 | 74.59±4.90 | Speed | | F_(2,40)_=963.41 | <0.001 | 0.980 |
|  |  |  |  |  |  |  | Phase | | F_(1,20)_=4.74 | 0.042 | 0.192 |
|  |  |  |  |  |  |  | Speed*Phase | | F_(2,40)_=7.48 | 0.002 | 0.272 |
| SW | 14.65±4.85 | 14.48±4.57 | 13.46±4.42 | 12.50±3.67 | 12.54±3.02 | 1.52±0.35 | Speed | | F_(2,40)_=7.71 | 0.001 | 0.278 |
|  |  |  |  |  |  |  | Phase | | F_(1,20)_=3.34 | 0.083 | 0.143 |
|  |  |  |  |  |  |  | Speed*Phase | | F_(2,40)_=1.70 | 0.196 | 0.078 |
| MoS | 2.96±1.40 | 3.27±1.46 | 3.60±1.19 | 3.50±1.07 | 3.48±0.94 | 12.30±2.52 | Speed | | F_(2,38)_=2.23 | 0.122 | 0.105 |
|  |  |  |  |  |  |  | Phase | | F_(1,19)_=0.06 | 0.811 | 0.003 |
|  |  |  |  |  |  |  | Speed*Phase | | F_(2,38)_=7.66 | 0.002 | 0.287 |
| SL_var_ | 2.57±0.99 | 2.67±0.85 | 1.60±0.28 | 1.76±0.45 | 1.45±0.42 | 2.70±0.61 | Speed | | F_(2,40)_=57.11 | <0.001 | 0.741 |
|  |  |  |  |  |  |  | Phase | | F_(1,20)_=2.09 | 0.164 | 0.095 |
|  |  |  |  |  |  |  | Speed*Phase | | F_(2,40)_=0.06 | 0.940 | 0.003 |
| SW_var_ | 1.58±0.25 | 1.88±0.56 | 1.87±0.41 | 2.29±0.50 | 2.34±0.43 | 3.31±0.96 | Speed | | F_(2,40)_=34.10 | <0.001 | 0.630 |
|  |  |  |  |  |  |  | Phase | | F_(1,20)_=25.81 | <0.001 | 0.563 |
|  |  |  |  |  |  |  | Speed*Phase | | F_(2,40)_=0.54 | 0.586 | 0.026 |
| MoS_var_ | 0.42±0.09 | 0.57±0.22 | 0.49±0.09 | 0.60±0.15 | 0.63±0.13 | 0.84±0.19 | Speed | | F_(2,38)_=70.79 | <0.001 | 0.788 |
|  |  |  |  |  |  |  | Phase | | F_(1,19)_=39.30 | <0.001 | 0.674 |
|  |  |  |  |  |  |  | Speed*Phase | | F_(2,38)_=0.68 | 0.511 | 0.035 |

All data are presented as mean ± SD in centimeters. BL: baseline, PP: post perturbation, SL: step length, SW: step width, MoS: mediolateral margin of stability, SL_var_: step length variability, SW_var_: step width variability, MoS_var_: mediolateral margin of stability variability.
